# Therapy to Teratology: Chronic Paternal Antioxidant Supplementation Alters Offspring Placental Architecture and Craniofacial Morphogenesis in a Mouse Model

**DOI:** 10.1101/2025.09.05.674556

**Authors:** Destani D. Derrico, Katherine Z. Scaturro, Erin E. Murray, Eliezar Guillen, Nathan S. Truss, Katherine A. Fairly, Samantha Higgins, Sanat S. Bhadsavle, Michael C. Golding

## Abstract

Oxidative stress plays a significant role in regulating the mammalian epigenome, with emerging evidence suggesting imbalances in the cellular redox state trigger stress-responsive epigenetic modifications that drive various human diseases. However, it remains unclear whether, like worms, epigenetic changes caused by redox imbalance or mitochondrial stress can move through the mammalian germline, potentially affecting the health of future generations. Antioxidant therapies are commonly used to reduce oxidative damage and are widely employed in cases of male infertility, where high-dose supplementation is often recommended to enhance sperm quality and overall measures of male reproductive health. Interestingly, in non-stressed, ostensibly healthy males, recent research suggests that antioxidants may have a negative influence on sperm epigenetic markers, indicating a potential epigenetic liability. However, whether male antioxidant treatment can induce paternal effects on offspring growth and development remains unknown. Here, we employed micro-CT imaging and geometric morphometrics to determine whether chronic antioxidant supplementation in healthy male mice affects placental growth and craniofacial development in their offspring. Adult C57BL/6J male mice were given a six-week preconception regimen of N-acetyl-L-cysteine (NAC; 400 mg/kg/day) and selenium (0.04 mg/kg/day), which continued throughout breeding with treatment-naïve females. Although we observed modest alterations to the histological patterning of the female placenta, placental weights and efficiency remained unchanged. In contrast, we observed significant changes in facial shape and symmetry in both male and female offspring, with female offspring exhibiting significant reductions in eye spacing and head area. These changes occurred without any macro changes in paternal metabolic health, indicating that antioxidant-induced shifts in redox balance may disrupt developmental programming in the male germline independent of changes in overall health. Our findings emphasize the need for caution when using antioxidants as preconception interventions and broadly suggest that modulation of the paternal redox axis may result in altered developmental programming and teratogenic effects.

## 1. Introduction

Oxidative stress is a potent regulator of the epigenetic landscape, influencing DNA methylation, histone modifications, and noncoding RNAs through both direct and indirect mechanisms.(1) These stress-induced epigenetic alterations, in turn, profoundly affect cellular function and have been implicated in the development of a wide range of human diseases, including age-related pathologies, neurodegenerative disorders, cancer, cardiovascular disease, infertility, and developmental disorders, including fetal alcohol spectrum disorders (FASDs).(1,2) However, a critical yet emerging area of investigation is the extent to which systemic oxidative stress is sensed by the cells of the reproductive tract and whether mitochondrially driven epigenetic changes can transfer through the mammalian germline to influence offspring health.

Emerging research in a wide range of lower-order sexually reproducing organisms suggests that epigenetic mechanisms can mediate the inter- and transgenerational inheritance of mitochondrial traits, revealing a potential pathway for oxidative stress-induced signaling across generations.(3) For example, in *C. elegans*, early-life mitochondrial stress induces adaptive changes in mitochondrial function that persist into adulthood and are transmitted *via* epigenetic mechanisms to subsequent generations, influencing offspring lifespan and stress resilience. Interestingly, these studies predominantly suggest that parental mitochondrial stress can induce heritable epigenetic changes in offspring that enhance mitochondrial function and confer increased stress resilience and longevity.(3)

In contrast to studies in worms, research using mammalian models demonstrates that dietary and environmental stressors promote the intergenerational transmission of adverse changes in mitochondrial function, resulting in detrimental impacts on offspring growth and long-term health. For example, studies examining rodents consistently demonstrate that a maternal low-protein or obesogenic diets negatively affect oocyte quality and disrupts the structure, function, and quality control of mitochondria.(4–7) These mitochondrial defects persist into fetal and postnatal life, resulting in lasting deficits in mitochondrial function, including compromised metabolism, increased oxidative stress, impaired mitophagy, and disrupted regulation of mitochondrial homeostasis, with lasting consequences on the health and stress resilience of the offspring.(8)

Similarly, work from our laboratory examining chronic paternal alcohol exposure demonstrates the paternal inheritance of adverse impacts on fetal development that strongly resemble the developmental defects observed in fetal alcohol spectrum disorders.(9) These impairments first appear during fetal development and are associated with altered placental growth and abnormalities in craniofacial development, including changes in facial shape and symmetry.(10–15) Notably, these developmental outcomes correlate with adverse effects on offspring mitochondrial function, including impaired complex I activity, shifts in the NADH/NAD^+^ ratio, and disrupted cellular metabolism that persist into adulthood and correlate with accelerated biological aging. Remarkably, paternal alcohol use appears to interact with maternal exposures to accelerate these mitochondrial deficits, promoting a sustained pro-inflammatory state, which increases offspring susceptibility to liver disease and hepatocellular carcinoma.(16–19)

Our ongoing research indicates that chronic paternal alcohol consumption impairs mitochondrial function within the male reproductive tract, correlating with alterations in sperm noncoding RNAs and long-lasting changes in offspring mitochondrial activity.(20,21) Given that reactive oxygen species activate multiple epigenetic and stress-response pathways that directly impact male fertility and epigenetic markers in sperm, (22) antioxidant interventions represent a logical intervention that could modify the epigenetic transmission of alcohol-related paternal effects.

Antioxidant therapies aim to mitigate oxidative stress, a condition characterized by an imbalance between reactive oxygen species and the cellular antioxidant defense.(23) While preclinical studies highlight the potential of antioxidants to mitigate cellular markers of oxidative damage, clinical outcomes have been inconsistent, with some reports even documenting adverse effects.(23,24) Indeed, in a recent study by Hug *et al*., modeling the effects of oxidative stress on the sperm epigenome, antioxidant therapy successfully corrected redox-induced epigenetic alterations. However, strikingly, non-stressed control animals exposed to the same antioxidants developed epigenetic changes of comparable magnitude to those caused by the original stressor.(25) These findings join a growing body of research suggesting that antioxidant treatments are not innocuous and, in the absence of oxidative stress, may disrupt normal epigenetic programming in the male germline.(26) However, whether these antioxidant-induced epimutations also transmit to offspring and affect developmental outcomes remains unknown. Therefore, as a preliminary step to investigating the ability of antioxidants to modify alcohol-induced paternal effects, we tested the hypothesis that a chronic antioxidant regimen would cause developmental changes in the offspring of non-stressed, antioxidant-treated fathers compared to unexposed controls.

N-acetyl-L-cysteine (NAC) and selenium (Se) are two widely studied antioxidant compounds used to investigate cellular responses to oxidative stress. NAC primarily functions as a precursor to cysteine, a rate-limiting amino acid required for the synthesis of glutathione (GSH), a major intracellular antioxidant. GSH neutralizes reactive oxygen species by donating electrons and converting to the oxidized glutathione disulfide (GSSG).(27) Similarly, selenium is a critical component of selenoproteins, including glutathione peroxidases (GPx), which catalyze the reduction of hydrogen peroxide and organic hydroperoxides using GSH as a substrate.

NAC and selenium supplementation have demonstrated beneficial effects on markers of male fertility, including specific epigenetic parameters.(28–30) Further, given that gestational NAC supplementation can attenuate the programmed susceptibility to obesity and insulin resistance in offspring of mothers maintained on a high-fat diet and correct FASD-related craniofacial phenotypes induced by gestational alcohol exposure (31,32), we selected these antioxidants for our studies. Our findings reveal that chronic male antioxidant treatment induces sex-specific effects on placental histological patterning and has an adverse impact on offspring craniofacial shape and morphology. These results suggest that disruptions to male redox balance by antioxidants, potentially as an unintended consequence of therapeutic use, may have teratogenic consequences on offspring development.

## 2. Materials and Methods

### 2.1 Ethics and Regulatory Compliance

We designed our study in accordance with the ARRIVE guidelines (33) and conducted all experiments in compliance with IACUC regulations and the National Research Council’s Guide for the Care and Use of Laboratory Animals, with prior approval from the Texas A&M University IACUC (protocol number 2023-0186).

### 2.2 Animal Studies and Antioxidant Exposures

We utilized male C57BL/6J strain mice (RRID:IMSR_JAX:000664) obtained from a breeder nucleus at the Texas A&M Institute for Genomic Medicine. We maintained males in the TIGM facility on a reverse 12-hour light/dark cycle (lights off at 8:30 AM) and fed them a standard chow diet (Catalog# 2019; Teklad Diets, Madison, WI, USA). Beginning on postnatal day 90, we individually housed each male to monitor the dosing of the antioxidant treatment. To help offset the stress of individual housing and minimize the impact of animal stress (34), we added shelter tubes (catalog# K3322; Bio-Serv, Flemington, NJ, United States) and additional nestlets to enhance cage enrichment, as described previously.(10,11,14)

We initiated the control and antioxidant treatments by exposing control mice to ultrafiltered water, while we exposed experimental mice to an antioxidant mixture comprised of 4.48 mg/mL N-acetyl-L-cysteine (catalog #A7250, Sigma-Aldrich, St. Louis, MO, United States) and 0.448 ug/mL selenium in the form of sodium selenite (catalog #S5261, Sigma-Aldrich, St. Louis, MO, United States). We selected these dosages from previous publications, anticipating maximum daily dosages of 400 mg/kg/day for NAC (32,35,36) and 0.04 mg/kg/day for selenium.(37) We added a zero-calorie flavor enhancer (0.0896% solution, Stevia in the Raw®, Cumberland Packing Corp., Brooklyn, NY, United States) to increase the palatability of the antioxidant treatment. Each week, we recorded the weight of each mouse (g) and the total weekly fluid consumption (g). We then quantified weekly fluid consumption by dividing the grams of fluid consumed by the sire’s body weight (g/g).

We maintained the preconception treatments for six weeks, which, in mice, encompasses approximately one complete spermatogenic cycle,(38) and continued treatments during the subsequent breeding phase. We paired control and antioxidant-treated males with naïve postnatal day 90 C57BL/6J strain dams, which we obtained from the Texas A&M Institute for Genomic Medicine (**Figure 1A**). We synchronized female reproductive cycles using the Whitten method (39), then placed one female in the male’s home cage. During this eight-hour breeding window, we substituted the antioxidant treatment with filtered water, ensuring females were not exposed to the antioxidant treatment. We confirmed matings by the presence of a vaginal plug, recorded female body weights, and returned females to their original cages. We rested males for two weeks, during which they continued the preconception control or antioxidant treatments, and then used them again in a subsequent mating. On gestational day ten, we confirmed pregnancy diagnosis by an increase in body weight of at least 1.8 g. We terminated dams on gestational day 16.5 using carbon dioxide asphyxiation followed by cervical dislocation, dissected the female reproductive tract, and recorded fetoplacental measures. We then collected digital photographs of the front, left, and right profiles of each fetus within each litter. We then either fixed the collected tissue samples in 10% neutral buffered formalin (catalog# 16004-128, VWR, Radnor, PA, United States) or snap-froze the tissues on dry ice and stored them at −80°C.

**Figure 1.**
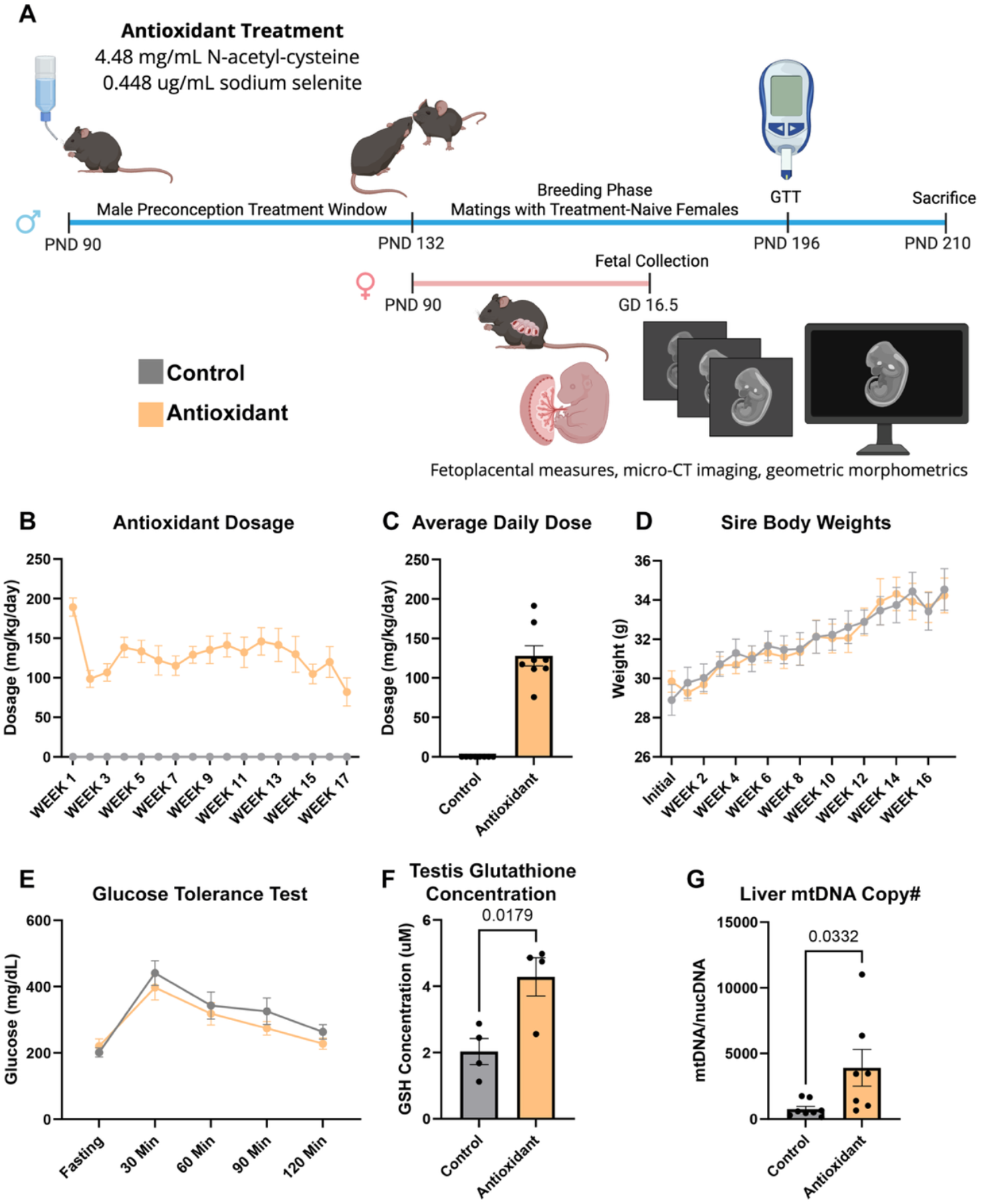
A mouse model to examine the paternal effects of chronic antioxidant supplementation on offspring development. **A**) Visual representation of the mouse model we used to determine the impacts of chronic paternal antioxidant supplementation on offspring growth and development. **B**) Average daily NAC dose (n=8) across the experimental course and **C**) cumulative average daily dose. **D**) Comparison of average weekly weight gain between sire treatment groups across the experimental course, including the 6-week preconception and 10-week breeding phases (n=8). **E**) Comparison of blood glucose levels during a glucose tolerance test. **F**) Measurement of testicular glutathione concentrations using a colorimetric assay (n=4). **G**) qPCR analysis of hepatic mitochondrial DNA copy number between treatments (n=8). We compared treatment groups using a two-way ANOVA and an unpaired Student’s t-test; data represent mean ± SEM.

### 2.3 Fetal sex determination

We isolated genomic DNA from the fetal tail using the HotSHOT method (40) and determined fetal sex using a PCR-based assay described previously (10).

### 2.4 Placental histological analysis

We examined the impact of preconception paternal antioxidant treatment on placental histology using previously described methods.(10,12,13) Briefly, we cut placentae in half and fixed one portion in neutral buffered formalin. We then randomly selected placentae across all litters, stained these samples with phosphotungstic acid to enhance tissue contrast, (41) and then processed them for Micro-Computed Tomography (micro-CT) imaging. We used Aquasonic Clear Ultrasound Gel (Catalog# 03-08; Parker Labs, Fairfield, NJ, United States) to prevent tissue desiccation during scanning. We imaged the treated samples using a SCANCO vivaCT 40 (SCANCO Medical AG, Brüttisellen, Switzerland) with a 55 kVp voltage x-ray tube and an exposure of 29 µA, yielding an image voxel size of 0.0105 mm^3^and a resolution of 95.2381 pixels/mm. We then used the open-source medical image analysis software Horos (Version 3.3.6; Nibble Co. LLC, Annapolis, Maryland, United States; https://horosproject.org/) to quantify layer-specific volumes, as described previously.(42)

### 2.5 Digital image acquisition and processing

During dissections, we collected digital photographs of the front, left, and right profiles of each fetus within each litter. We then processed the images for morphometric analyses using methods described previously.(14,15) Briefly, we imported digital images of the facial profiles into the publicly available software **tpsUtil32** ((44); version 1.83) to generate TPS files for landmarking. We then used the publicly available software **tpsDig2w64** ((43) version 2.32) for image analysis by first setting the reference scale bar in the picture to 1 mm and then demarcating the eighteen facial landmarks described previously (14,15) for the front profile and twenty-two landmarks for the side profiles (**Table 1**). To ensure consistency, a single individual (N.S.T.) demarcated the landmarks in each photograph, consistently identifying the exact location and order for each image. To add additional landmarks, we generated the outline around the head using the publicly available program **tpsDig2w64** ((43) version 2.32), producing a total of 47 landmarks for the front profile and a total of 51 landmarks for the side profiles. In curating this dataset, we named each file with the litter I.D., sex, and uterine position of each fetus. Finally, we used the publicly available program **tpsUtil32** ((44); version 1.83) to create our final TPS files, inclusive of all landmarks for use in the MorphoJ software for analysis.

**Table 1.** Landmarks used in the morphometric analysis of the front, left, and right facial profiles.

| Number | Front View | Left/Right View |
| --- | --- | --- |
| 1 | Highest Head Point | Nose Tip |
| 2 | Bottom of Mandible | Nasion |
| 3 | Right Corner of Mouth | Highest Head Point |
| 4 | Left Corner of Mouth | Curve of Skull |
| 5 | Top of Philtrum | Back of Skull |
| 6 | Bottom of Philtrum | Bottom of Mandible |
| 7 | Nose Tip | Front of Mandible |
| 8 | Nasion | Upper Philtrum |
| 9 | 3 O'clock Eye Position (Left) | Inner Mouth |
| 10 | 12 O'clock Eye Position (Left) | Snout Edge |
| 11 | 9 O'clock Eye Position (Left) | Central Auditory Canal |
| 12 | 6 O'clock Eye Position (Left) | 3 O'clock Eye Position |
| 13 | Pupil Center (Left) | 12 O'clock Eye Position |
| 14 | 3 O'clock Eye Position (Right) | 9 O'clock Eye Position |
| 15 | 12 O'clock Eye Position (Right) | 6 O'clock Eye Position |
| 16 | 9 O'clock Eye Position (Right) | Pupil Center |
| 17 | 6 O'clock Eye Position (Right) |  |
| 18 | Pupil Center (Right) |  |

### 2.6 Geometric Morphometrics and Statistical Analyses of Facial Images

We imported the generated TPS. files for each fetus into the MorphoJ software ((45) version build 1.07a, Java version 1.8.0_291 (Oracle Corporation)) and conducted geometric morphometric analysis using methods described previously.(14,15) Briefly, we added classifiers describing each treatment group and then separately normalized the datasets for scale, rotation, and translation using the Procrustes fit feature (45). We then generated a covariance matrix, which we used to conduct Principal Component Analysis (PCA).

We then used Canonical Variate (CV) analysis to identify differences in facial features between treatments and exported the raw CV scores into the publicly available Paleontological Statistics Software Package for Education and Data Analysis (PAST) analysis software ((46) version 4.03; [https://softfamous.com/postdownload-file/past/18233/13091/.]). We conducted multivariate analyses of the raw CV scores using statistical methods described previously.(47–50) These included the parametric Multivariate analysis of variance (MANOVA), and Nonparametric Analysis of similarities (ANOSIM), and Permutational multivariate analysis of variance (PERMANOVA) tests, followed by Bonferroni correction. We generated the CV lollipop and scatter plots using the graphing features of MorphoJ (45).

### 1.8 Data handling and statistical analysis

We subjected all data generated during this study to the data management practices and statistical analyses described previously.(18,19) Briefly, we recorded our initial observations by hand and then inserted these measurements into Google Sheets or Microsoft Excel. In line with modern statistical reporting,(51) we have moved away from binary significance labels and now interpret p-values as graded evidence against the null hypothesis. We consider p-values below 0.01 to be strong evidence for an effect, while p-values between 0.1 and 0.01 provide moderate evidence of an effect (52). Here, we report the exact p-values for each test.

We transferred the collected datasets into GraphPad Prism 10 (RRID:SCR_002798, GraphPad Software Inc., La Jolla, CA, USA). We first employed the ROUT test (Q = 1%) to identify outliers and verified equal variance using either the Brown-Forsythe or F testing. If data passed normality and variance testing (alpha = 0.05), we employed either an unpaired, parametric (two-tailed) t-test or a One-way or Two-way ANOVA. We then used Šídák’s multiple comparisons test or Tukey’s post hoc test to compare each treatment to the control. If, however, the collected datasets failed the test for normality or we observed unequal variance, we ran a Kruskal-Wallis test followed by Dunn’s multiple comparisons test. We present detailed descriptions of each statistical test and the sample sizes for each figure in **Table 2**.

**Table 2.** Statistical Tests Employed In Each Figure.

| Figure | Sample Size | # Outliers | Analysis |
| --- | --- | --- | --- |
| 1B | CC: 8<br>CAX: 8 | CC: 0<br>CAX: 0 | Two-way ANOVA:<br>Šídák's multiple<br>comparisons test |
| 1C | CC: 8<br>CAX: 8 | CC: 0<br>CAX: 0 | Unpaired t-test |
| 1D | CC: 8<br>CAX: 8 | CC: 0<br>CAX: 0 | Mixed-effects analysis:<br>Dunnett's multiple<br>comparisons test |
| 1E | CC: 8<br>CAX: 8 | CC: 0<br>CAX: 0 | AUC, Two-way ANOVA:<br>Dunnett's multiple<br>comparisons test |
| F | CC: 8<br>CAX: 8 | CC: 0<br>CAX: 0 | Unpaired t-test |
| G | CC: 8<br>CAX: 8 | CC: 0<br>CAX: 0 | Unpaired t-test |
| 2A - D | CC: 14 males, 20 females; 4 litters<br>CAX: 22 males, 21 females; 5 litters | CC: 0<br>CAX: 0 | Linear mixed model,<br>Mixed-effects analysis:<br>Šídák's multiple<br>comparisons test |
| 2E - J | CC: 9 males, 8 females; 4 litters<br>CAX: 12 males, 12 females; 5 litters | CC: 0<br>CAX: 0 | Two-way ANOVA:<br>Tukey's multiple<br>comparisons test |
| 3B, D | CC: 12 males, 12 females; 3 litters<br>CAX: 20 males, 17 females; 4 litters | CC: 0<br>CAX: 0 | Canonical variate analysis |
| 3E | CC: 12 males, 12 females; 3 litters<br>CAX: 20 males, 17 females; 4 litters | CC: 0<br>CAX: 0 | Procrustes ANOVA |
| 4A - J | CC: 12 males, 12 females; 3 litters<br>CAX: 20 males, 17 females; 4 litters | CC: 0<br>CAX: 0 | Two-way ANOVA:<br>Tukey's multiple<br>comparisons test |

## 3. Results

### A mouse model to examine the impacts of chronic antioxidant supplementation on offspring fetoplacental growth and craniofacial development

To investigate the effects of chronic paternal antioxidant supplementation on offspring development, we exposed males to the preconception treatments for six weeks and then bred the exposed males with treatment-naïve dams (**Figure 1A**). Over the six-week preconception and subsequent ten-week breeding phase, exposed males received an average daily dose of 127 g/kg/day NAC (**Figure 1B-C**) and 0.013 mg/kg/day selenium (data not shown). We did not identify any differences in sire body weights between the control and antioxidant treatment groups (**Figure 1D**).

In rodent models examining type 2 diabetes or diet-induced metabolic syndrome, NAC often improves glucose tolerance and reduces fasting blood glucose levels.(53) However, we did not identify any impacts of the paternal antioxidant treatment on sire fasting blood glucose levels or during glucose tolerance testing (**Figure 1E**). Similarly, Dual-Energy X-ray Absorptiometry (DEXA) scanning did not identify any impacts of the antioxidant treatment on body fat percentage (data not shown).

We next assessed sire glutathione concentrations. As anticipated (27), NAC treatment increased cellular glutathione levels, including a 50% increase in the testis (*p*= 0.0179, **Figure 1F**). Hepatic mitochondrial DNA (mtDNA) copy number serves as a crude proxy of systemic mitochondrial stress.(54) In previous studies, NAC treatment ameliorated alcohol-induced increases in hepatic mitochondrial stress, including increases in mtDNA.(55) Unexpectedly, we identified modest evidence of increased mitochondrial stress in antioxidant-treated males, which exhibited an 80% increase in mtDNA copy number compared to controls (**Figure 1G**).

### Preconception male antioxidant supplementation modifies female placental histological patterning

After six weeks of treatment, corresponding to approximately one complete spermatogenic cycle, we bred treated males with treatment-naïve females. Of the eight exposed males in each treatment, four control males and five antioxidant-treated males sired litters, which we used in our comparisons. After diagnosing pregnancy on gestational day ten, we ceased all animal handling and left dams undisturbed until gestational day 16.5 (GD16.5). We then sacrificed pregnant dams, excised the female reproductive tract, and collected multiple measures of offspring fetoplacental growth. As in our previous studies, we selected GD16.5, as this time point represents the phase of pregnancy during which placental growth (in terms of diameter, thickness, and weight) has plateaued, while fetal growth continues to increase.(56,57)

We first used a linear mixed model to compare measures of fetoplacental growth between treatments and then followed these analyses with a two-way ANOVA to contrast the effects of the preconception antioxidant treatments and offspring sex. We observed a modest 5% decline in female offspring body weight (*p*= 0.1002), but no differences in male offspring (**Figure 2A**). We did not observe any effects of paternal antioxidant supplementation on crown-rump lengths, placental weights, or placental efficiency in either male or female offspring (**Figure 2B-D**). We did not observe any treatment effects on gestation length, litter size, or offspring sex ratio among the experimental treatments (data not shown).

**Figure 2.**
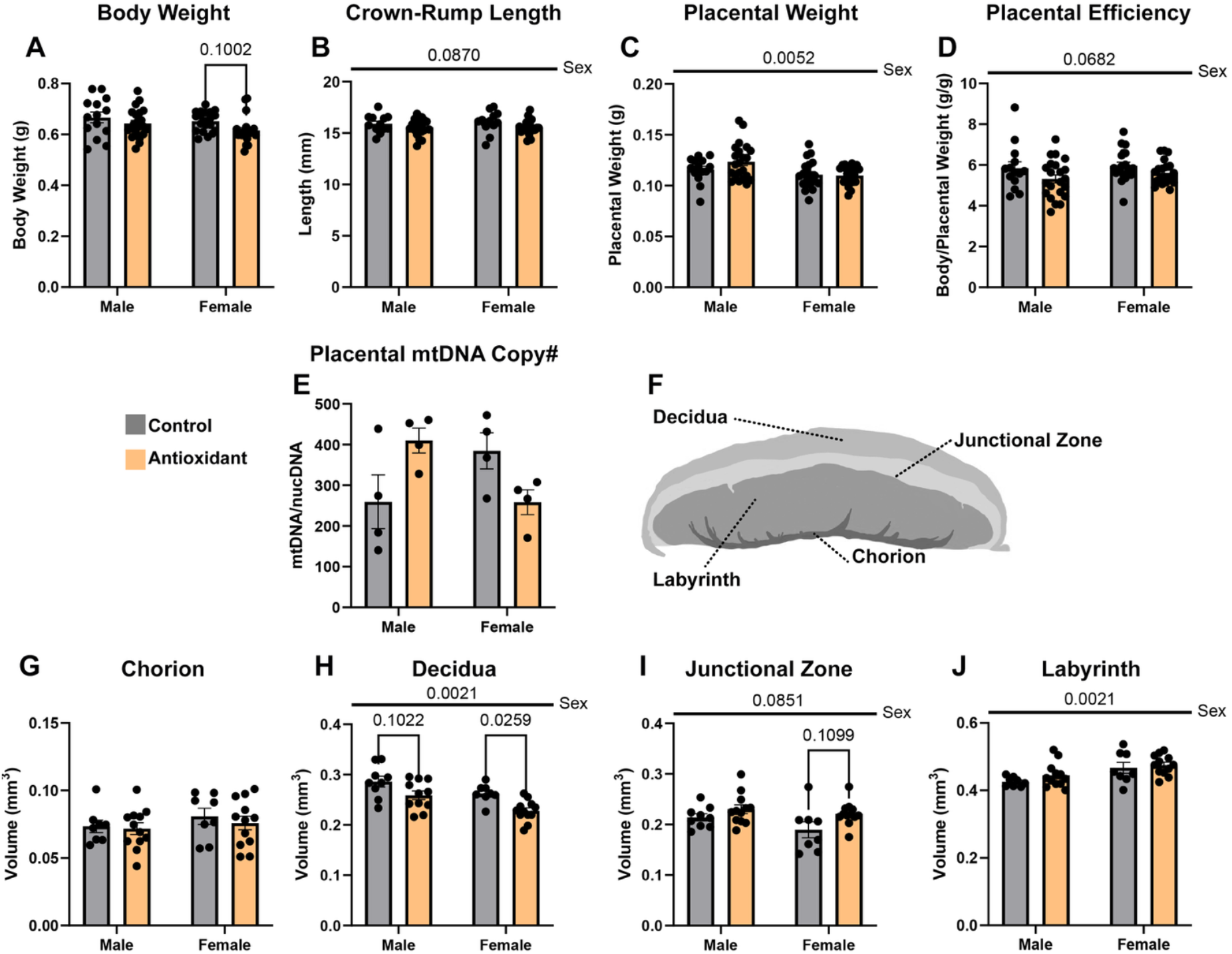
Chronic high-dose antioxidant supplementation modifies offspring placental histological patterning. We used a linear mixed model with repeated measures, followed by a two-way ANOVA, to compare the combined effects of preconception paternal antioxidant treatment on male and female offspring (**A**) fetal weights, (**B**) crown-rump lengths, (**C**) placental weights, and (**D**) placental efficiency (n = 4 to 5 litters). (**E**) We used qPCR to compare placental mitochondrial DNA copy number in male and female placentae derived from each treatment group (n = 4). (**F**) Schematic diagram depicting the histological layers of the murine placenta. Using phosphotungstic acid staining to enhance tissue contrast and micro-CT imaging, we compared the proportional volumes of the placental (**G**) chorion, (**H**) decidua, (**I**) junctional zone, and (**J**) labyrinth in male and female offspring sired by males from the control and antioxidant preconception treatment groups. We used a two-way ANOVA followed by either Sidak’s or Tukey’s post-hoc testing to compare treatment and sex; data represent mean ± SEM.

As a rough measure of mitochondrial health, we used quantitative PCR (qPCR) to assay mitochondrial DNA (mtDNA) copy number in the placenta. We did not observe any differences in mtDNA copy number in either male or female placentae (**Figure 2E**). We next used micro-CT imaging to determine the impacts of chronic paternal antioxidant exposure on placental patterning and histological organization. This technique enables the three-dimensional quantification of the murine placenta, allowing for discrimination and proportional comparisons of the placental chorion, labyrinth, junctional zone, and decidua layers (**Figure 2F**).(41,42) This analysis revealed ∼10% to 15% decreases in the proportional volume of male and female decidua, respectively (*p*= 0.1022 and *p*= 0.0259), and a 15% increase in the proportional volume of the female junctional zone (*p*= 0.1099). We did not identify any differences in the proportional volume of the placental chorion or labyrinth layers (**Figure 2G-J**).

### Preconceptional male antioxidant supplementation modifies offspring craniofacial shape and symmetry

To determine the effects of chronic paternal antioxidant supplementation on craniofacial growth and patterning, we employed geometric morphometrics, a widely used technique for comparing diverse aspects of craniofacial shape, including the characterization of fetal alcohol syndrome-associated facial dysmorphology in both preclinical and clinical studies.(58–60) Using digital images collected during dissections, we utilized MorphoJ to perform a generalized Procrustes analysis, removing scale, orientation, and rotation from the dataset and placing it within a common coordinate system. We then performed a Procrustes ANOVA to quantify relative variations in the proportional size and shape through the positions of facial landmarks attributable to one or more factors in our model. We identified strong evidence of changes in the shape of the left and front facial profiles, but did not identify any changes in the right profile (**Supplemental Table S2**). We did not identify evidence of differences in centroid size for any of the profiles.

We then conducted Canonical variate (CV) analysis, which identified alterations in the growth of the jaw and positioning of the eyes and ears (**Figure 3A**). CV analysis of geometric facial relationships in the left profile revealed paternal antioxidant supplementation induced a morphometric shift away from the control treatment along canonical variate one and to a lesser extent, canonical variate two, which together accounted for approximately 45% and 41% of the observed variance in our model (**Figure 3B**). Similarly, analysis of the front profile revealed shifts in canonical variates one and two, with a shift of midline features to the left (**Figure 3C-D**).

**Figure 3.**
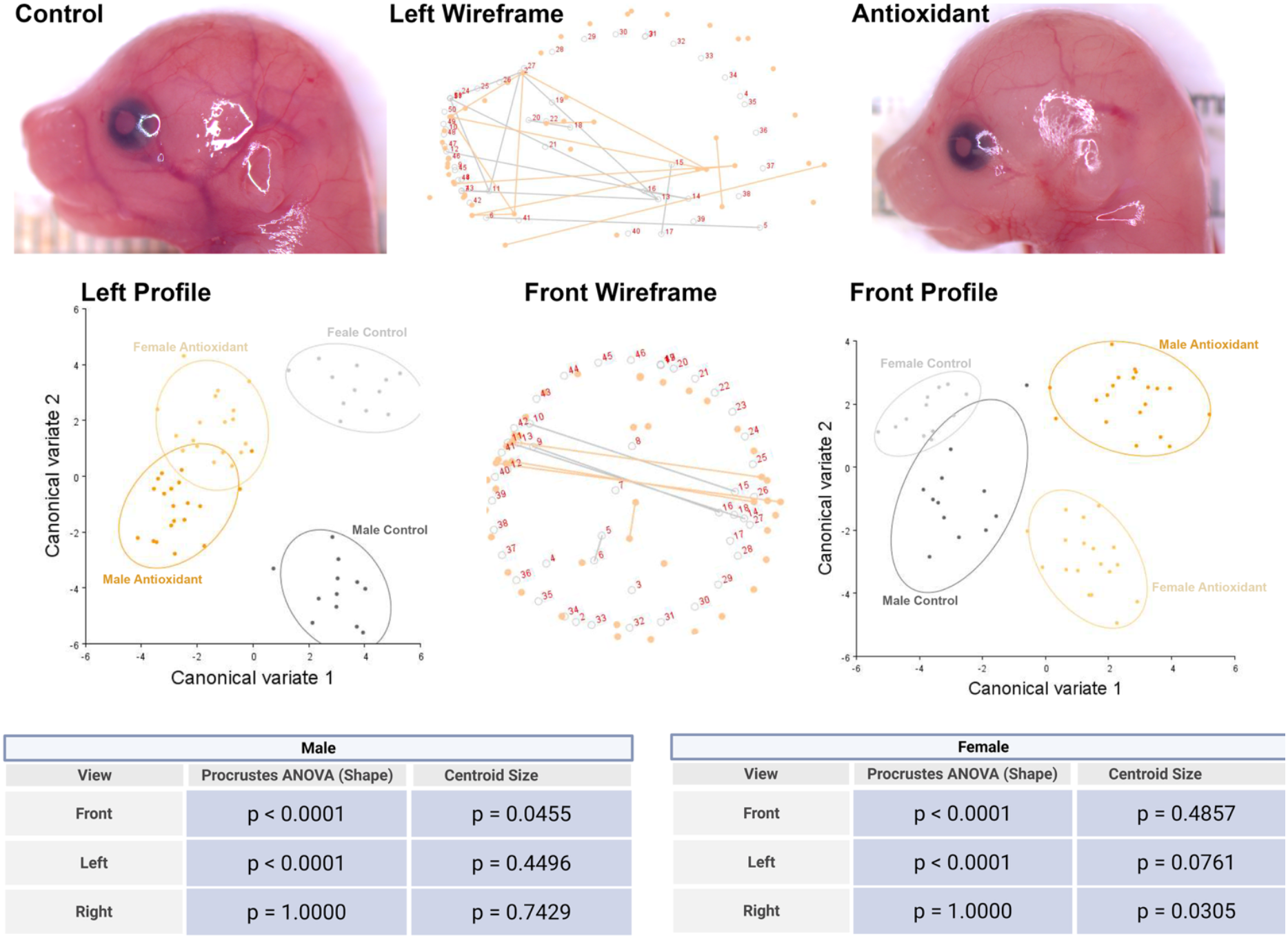
Preconception paternal antioxidant supplementation alters offspring craniofacial shape and symmetry. We employed geometric morphometrics and Procrustes analysis of variance (ANOVA), followed by canonical variate (CV) analysis, to evaluate the effects of paternal antioxidant treatment on facial shape and symmetry. (**A**) Representative images of male offspring derived from control (left) and antioxidant-treated males (right) fetuses flanking a wireframe graph of CV1 (center), illustrating the relative shifts in facial landmarks in male offspring. (**B**) CV plot depicting treatment-induced changes in the left facial profile. (**C**) Wireframe graph of CV1 and (**D**) CV plot depicting treatment-induced changes in the front facial profile. We used Procrustes ANOVA to evaluate the effects of paternal antioxidant supplementation on the facial shape and symmetry of (**E**) male and female offspring.

We then used the raw CV scores to conduct three independent multivariate analyses, including MANOVA, ANOSIM, and PERMANOVA, followed by Bonferroni correction to identify significant differences in clustering and distance between treatment groups. Each of these statistical tests revealed strong evidence (*p*< 0.0006) of pairwise differences between the treatment groups (**Supplemental Table S1**). We then separated males and females and performed a Procrustes ANOVA. These analyses revealed an impact of paternal antioxidant supplementation on the shape of the left and front profiles of male and female offspring but not the right (**Figure 3E**). This analysis also identified changes in centroid size for the male front profile and female left profile, indicating a shift in facial size independent of shape (**Figure 3E**). Therefore, paternal antioxidant treatment induced changes in facial shape and allometry.

### Preconceptional male antioxidant supplementation induces sex-specific changes in eye spacing and head size

To further explore the effects of paternal antioxidant treatments on craniofacial growth and development, we compared linear measures utilizing established facial measurements routinely employed in studies examining mouse models of craniofacial dysgenesis (**Figure 4A**).(61) We observed strong evidence for an effect of paternal antioxidant supplementation on female outer canthal distance, with the offspring of antioxidant-exposed fathers displaying a 6.5% reduction (*p*= 0.0010, **Figure 4B**). We also identified a 4% reduction in female interpupillary distance (*p*= 0.0010) but did not identify evidence to support effects on inner canthal distance or philtrum length (**Figure 4C-E**).

**Figure 4.**
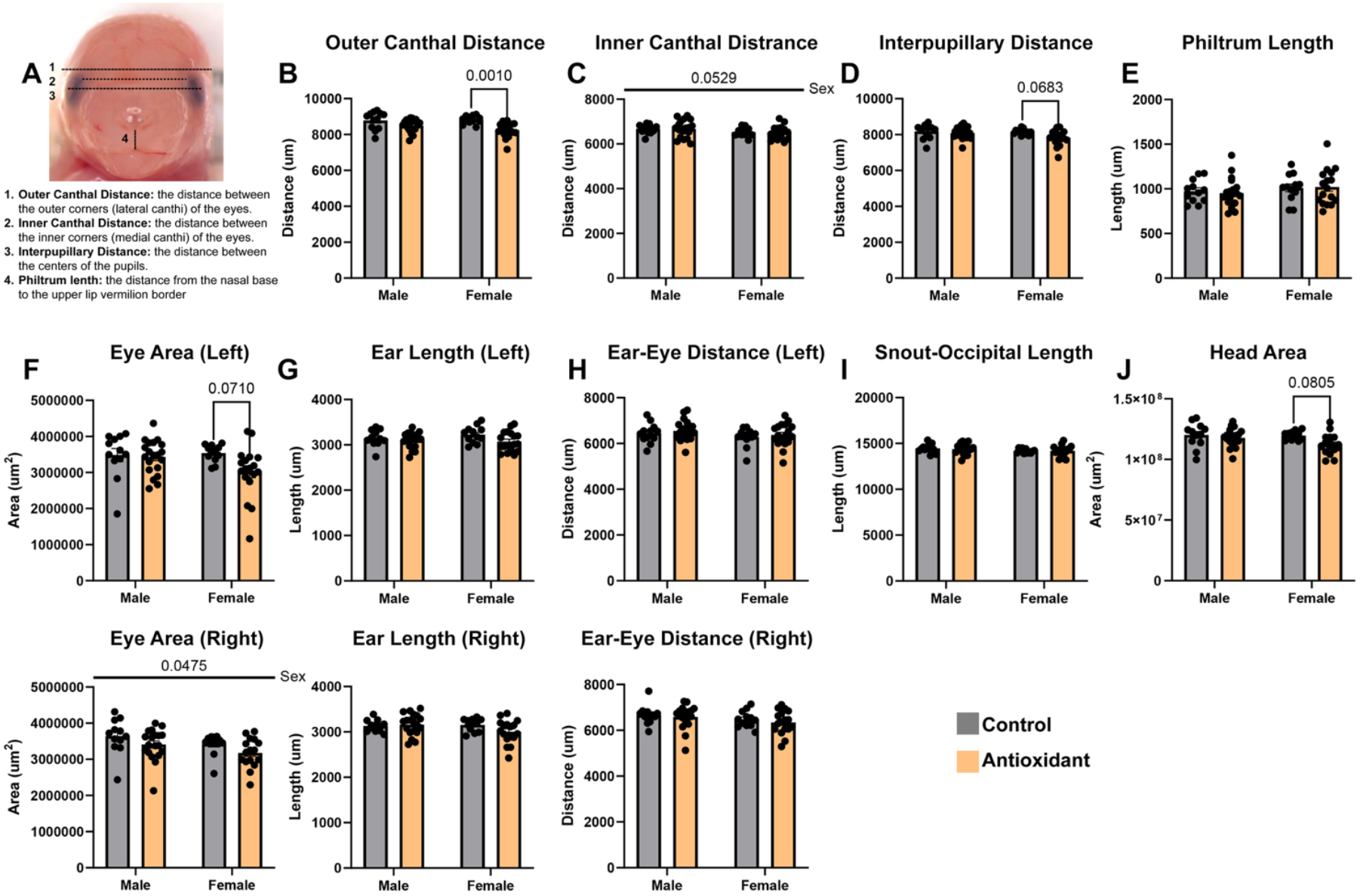
Preconception paternal antioxidant supplementation exerts sex-specific changes in linear measurements of offspring eye spacing and head size. (**A**) Graphic representation of the employed measures of craniofacial morphology in the frontal profile, which are established measures disrupted in mouse models examining prenatal alcohol exposure.(61) We used a two-way ANOVA followed by Tukey’s post-hoc testing to determine the effects of the paternal antioxidant treatment on (**B**) outer canthal distance, (**C**) inner canthal distance, (**D**) interpupillary distance, (**E**) philtrum length (**F**) eye area, (**G**) ear length, (**H**) ear-eye distance, (**I**) snout-occipital length, and (**J**) head area. Data represent mean ± SEM.

In the female offspring of antioxidant-treated males, we identified a 15% reduction in the area of the left eye but did not identify any differences in the right (*p*= 0.0710, **Figure 4F**). We did not identify any differences in the left or right ear length, or ear-eye distance in either male or female offspring (**Figure 4G-H**). We did not identify any differences in snout-occipital length for either male or female offspring, but did identify a 6% decrease in head area for female offspring (*p*= 0.0805, **Figure 4I-J**). Collectively, these studies reveal that paternal antioxidant supplementation alters the size and spacing of the eyes and head area in female offspring.

## 4. Discussion

Our previous studies demonstrate that the offspring of alcohol-exposed males share a signature of hepatic mitochondrial stress and redox imbalance that resembles the metabolic disturbances observed in actively drinking males.(62,63) These observations suggest that the paternal germline may sense this toxicant-induced redox imbalance and modulate similar pathways in the early embryo, with lasting impacts on development and metabolic function.(16–19) Based on these observations, we speculated that antioxidant supplementation may ameliorate the alcohol-induced paternal effects on placental and craniofacial growth observed in our model. However, recent studies strongly indicate that antioxidant supplementation alone modifies the sperm epigenome, suggesting disruptions in male redox health, generally, may induce paternal effects on offspring development (*reviewed* (26)). Herein, we sought to determine if a chronic antioxidant regimen could induce paternal effects on offspring fetoplacental growth and craniofacial development. Our findings reveal that chronic paternal antioxidant treatment induced sex-specific effects on offspring histological patterning. More strikingly, our work also reveals that administration of antioxidants to ostensibly non-stressed males alters craniofacial shape and symmetry in both male and female offspring, suggesting that chronic redox interventions themselves carry an intergenerational liability.

N-acetylcysteine (NAC) is a widely used antioxidant with therapeutic applications across multiple health conditions. It is also commonly taken as a dietary supplement to support athletic performance, cognitive function, and general wellness.(64) In addition to serving as a precursor for glutathione synthesis, NAC can activate the endogenous antioxidant response by modifying cysteine residues on Kelch-like ECH-associated protein 1 (KEAP1). This modification disrupts KEAP1’s repression of NRF2, allowing NRF2 to translocate to the nucleus and induce the transcription of a broad array of cytoprotective genes, including those encoding antioxidant enzymes.(65) Due to these dual mechanisms of action, high-dose NAC is used as a frontline treatment for acetaminophen overdose, as a therapeutic agent in congestive heart failure, and as a recovery aid following intense physical exertion. It is also popular among self-described biohackers seeking to counteract reactive oxygen species, which are implicated in aging and chronic disease.(64,66–68) In toxicology research, NAC is frequently used as a protective agent, including in studies examining alcohol-induced craniofacial birth defects, where it is believed to buffer against teratogenic effects.(32)

Despite its broad use, a growing body of evidence indicates that NAC may disrupt normal physiology in non-stressed systems. For instance, in otherwise healthy mice, NAC supplementation reduced mitochondrial activity in brown adipose tissue and increased markers of oxidative stress within mitochondria.(69) In another study, NAC-induced reductive stress, impaired insulin signaling, and glucose transport in muscle and adipose cells of normoglycemic mice, while paradoxically improving both outcomes in diabetic animals.(70) These findings suggest that NAC supplementation as a preventive or baseline treatment in non-stressed systems may not be universally beneficial. These preclinical observations align with broader evidence that antioxidant overuse can attenuate physiological adaptations, including those triggered by endurance training, such as increased mitochondrial biogenesis, enhanced cellular defense mechanisms, and improved insulin sensitivity.(71,72)

Broadly, the term ‘antioxidant’ describes a wide range of compounds that either directly scavenge reactive oxygen species or bolster the biological pathways that facilitate ROS detoxification. While antioxidants have therapeutic potential, particularly in the context of oxidative stress-related disorders, this term has also become highly marketable. This interest derives from widespread public awareness of free radical damage and its links to aging and chronic disease. Consequently, supplement manufacturers frequently emphasize the antioxidant content of their products, often by including standard vitamins such as C and E or by branding products with labels like “antioxidant blend” or “antioxidant support.” However, emerging research has shown that high doses of certain antioxidants, including N-acetylcysteine or vitamin E, suppress the normal physiological ROS underpinning normal cellular signaling, shifting the cellular redox environment toward a reductive state, which is itself pathological. Consequently, while intended to promote health, the indiscriminate use of antioxidant supplements may paradoxically impair cellular function, particularly in otherwise healthy individuals seeking to enhance fertility or overall well-being. Importantly, these compounds are not biologically inert, and their effects, particularly when delivered in combination, are very poorly understood.

While this preliminary study produced several compelling observations, it is important to acknowledge several limitations. First, we cannot definitively determine whether the antioxidant treatment directly affected sperm production and maturation or if the observed paternal effect is a downstream consequence of mitochondrial stress in a distant tissue, such as the liver. Second, although our discussion primarily focused on NAC, we utilized an antioxidant cocktail comprised of both NAC and selenium. Additionally, we included a low dose of the commercial sweetener stevia to enhance palatability, which itself may possess antioxidant properties.(73) Therefore, we cannot conclusively determine if the observed antioxidant effects resulted from a single component or a synergistic effect of the combination. Our primary goal, however, was to employ a treatment with the maximal chance of modifying phenotypes that shift in response to chronic paternal alcohol use. Third, our study utilized a C57BL/6J murine model. While highly informative, this strain is known to be sensitive to redox stress and may not fully recapitulate human reproductive biology. Furthermore, our antioxidant treatment was delivered systemically and chronically, a regimen that may differ from the intermittent or more targeted approaches commonly employed in clinical settings. Further, our outcome measures focused predominantly on fetal and placental metrics; therefore, the long-term consequences for postnatal health and aging remain unexplored and warrant further investigation in future studies. Finally, although we presume that the observed paternal effects are transmitted *via* epigenetic mechanisms, we do not assay any epigenetic measures, including DNA methylation, chromatin organization, or noncoding RNAs. Future studies will compare epigenetic changes in sperm induced by antioxidants to those we observe in alcohol-exposed sperm.

## Conclusions

Although studies in *C. elegans* have shown that mitochondrial stress in the F0 generation can influence bioenergetic function across subsequent generations, evidence for transmissible mitochondrial effects in mammals remains limited.(3) However, emerging research suggests that similar pathways may operate in mammals, where mitochondrial toxicants or agents that alter the redox environment could have heritable effects on offspring development and health. Our findings extend teratogenic concerns beyond established mitochondrial toxicants like alcohol to include a broader range of environmental exposures and dietary supplements that disrupt redox balance. These results highlight the need for comprehensive preconception counseling for both parents and support expanding epidemiological studies to examine not only impacts on sperm count and fertility but also long-term developmental outcomes in offspring.

## Supplementary Materials

Supplemental Table S1: Detailed descriptions of the statistical tests and the sample sizes employed in our study.

Supplemental Table S2: MorphoJ Analysis of Offspring Craniofacial Shape.

## Author Contributions

Conceptualization, DDD, and MCG. Methodology, DDD, KZS, and MCG. Investigation, DDD, KZS, EEM, EG, NST, KAF, SH, SSB, and MCG, Formal analysis, Visualization DDD, KZS, SSB, and MCG. Funding acquisition, Supervision, MCG. Writing, DDD, and MCG.

## Institutional Review Board Statement

The study was conducted with prior approval from the Texas A&M University IACUC (protocol number 2023-0186).

## Data Availability Statement

The raw data supporting the conclusions of this article will be made available by the authors on request.

## Citations

1. Kreuz S, Fischle W. Oxidative stress signaling to chromatin in health and disease. Epigenomics. 2016 June;8(6):843–62.

2. Terracina S, Tarani L, Ceccanti M, Vitali M, Francati S, Lucarelli M, et al. The Impact of Oxidative Stress on the Epigenetics of Fetal Alcohol Spectrum Disorders. Antioxidants (Basel). 2024 Mar 28;13(4):410.

3. Zhang Q, Tian Y. Molecular insights into the transgenerational inheritance of stress memory. J Genet Genomics. 2022 Feb;49(2):89–95.

4. Mitchell M, Schulz SL, Armstrong DT, Lane M. Metabolic and Mitochondrial Dysfunction in Early Mouse Embryos Following Maternal Dietary Protein Intervention. Biol Reprod. 2009 Apr;80(4):622–30.

5. Saben JL, Boudoures AL, Asghar Z, Thompson A, Drury A, Zhang W, et al. Maternal Metabolic Syndrome Programs Mitochondrial Dysfunction via Germline Changes across Three Generations. Cell Rep. 2016 June 28;16(1):1–8.

6. Ferey JLA, Boudoures AL, Reid M, Drury A, Scheaffer S, Modi Z, et al. A maternal high-fat, high-sucrose diet induces transgenerational cardiac mitochondrial dysfunction independently of maternal mitochondrial inheritance. Am J Physiol Heart Circ Physiol. 2019 May 1;316(5):H1202–10.

7. Marei WFA, Smits A, Mohey-Elsaeed O, Pintelon I, Ginneberge D, Bols PEJ, et al. Differential effects of high fat diet-induced obesity on oocyte mitochondrial functions in inbred and outbred mice. Sci Rep. 2020 June 17;10(1):9806.

8. Di Berardino C, Peserico A, Capacchietti G, Zappacosta A, Bernabò N, Russo V, et al. High-Fat Diet and Female Fertility across Lifespan: A Comparative Lesson from Mammal Models. Nutrients. 2022 Oct 17;14(20):4341.

9. Golding MC. Teratogenesis and the epigenetic programming of congenital defects: Why paternal exposures matter. Birth Defects Res. 2023 July 9;

10. Thomas KN, Zimmel KN, Roach AN, Basel A, Mehta NA, Bedi YS, et al. Maternal background alters the penetrance of growth phenotypes and sex-specific placental adaptation of offspring sired by alcohol-exposed males. FASEB J. 2021 Dec;35(12):e22035.

11. Thomas KN, Zimmel KN, Basel A, Roach AN, Mehta NA, Thomas KR, et al. Paternal alcohol exposures program intergenerational hormetic effects on offspring fetoplacental growth. Front Cell Dev Biol. 2022;10:930375.

12. Roach AN, Zimmel KN, Thomas KN, Basel A, Bhadsavle SS, Golding MC. Preconception paternal alcohol exposure decreases IVF embryo survival and pregnancy success rates in a mouse model. Mol Hum Reprod. 2023 Jan 31;29(2):gaad002.

13. Bhadsavle SS, Scaturro KZ, Golding MC. Maternal 129S1/SvImJ background attenuates the placental phenotypes induced by chronic paternal alcohol exposure. Reprod Toxicol. 2024 June;126:108605.

14. Thomas KN, Srikanth N, Bhadsavle SS, Thomas KR, Zimmel KN, Basel A, et al. Preconception paternal ethanol exposures induce alcohol-related craniofacial growth deficiencies in fetal offspring. J Clin Invest. 2023 June 1;133(11):e167624.

15. Higgins SL, Bhadsavle SS, Gaytan MN, Thomas KN, Golding MC. Chronic paternal alcohol exposures induce dose-dependent changes in offspring craniofacial shape and symmetry. Front Cell Dev Biol. 2024;12:1415653.

16. Chang RC, Wang H, Bedi Y, Golding MC. Preconception paternal alcohol exposure exerts sex-specific effects on offspring growth and long-term metabolic programming. Epigenetics Chromatin. 2019 Jan 22;12(1):9.

17. Chang RC, Thomas KN, Bedi YS, Golding MC. Programmed increases in LXRα induced by paternal alcohol use enhance offspring metabolic adaptation to high-fat diet induced obesity. Mol Metab. 2019 Dec;30:161–72.

18. Basel A, Bhadsavle SS, Scaturro KZ, Parkey GK, Gaytan MN, Patel JJ, et al. Parental Alcohol Exposures Associate with Lasting Mitochondrial Dysfunction and Accelerated Aging in a Mouse Model. Aging Dis. 2024 July 27;

19. Basel A, Bhadsavle SS, Scaturro KZ, Parkey GK, Jones-Hall Y, Golding MC. Parental Alcohol Use Disrupts Offspring Mitochondrial Activity, Promoting Susceptibility to Toxicant-Induced Liver Cancer. Aging Dis. 2025 Jan 31;

20. Bedi Y, Chang RC, Gibbs R, Clement TM, Golding MC. Alterations in sperm-inherited noncoding RNAs associate with late-term fetal growth restriction induced by preconception paternal alcohol use. Reprod Toxicol. 2019 Aug;87:11–20.

21. Roach AN, Bhadsavle SS, Higgins SL, Derrico DD, Basel A, Thomas KN, et al. Alterations in sperm RNAs persist after alcohol cessation and correlate with epididymal mitochondrial dysfunction. Andrology. 2023 Dec 3;

22. Bisht S, Faiq M, Tolahunase M, Dada R. Oxidative stress and male infertility. Nat Rev Urol. 2017 Aug;14(8):470–85.

23. Forman HJ, Zhang H. Targeting oxidative stress in disease: promise and limitations of antioxidant therapy. Nat Rev Drug Discov. 2021;20(9):689–709.

24. Al-Madhagi H, Masoud A. Limitations and Challenges of Antioxidant Therapy. Phytother Res. 2024 Dec;38(12):5549–66.

25. Hug E, Renaud Y, Guiton R, Ben Sassi M, Marcaillou C, Moazamian A, et al. Exploring the Epigenetic Landscape of Spermatozoa: Impact of Oxidative Stress and Antioxidant Supplementation on DNA Methylation and Hydroxymethylation. Antioxidants (Basel). 2024 Dec 12;13(12):1520.

26. Moazamian A, Saez F, Drevet JR, Aitken RJ, Gharagozloo P. Redox-Driven Epigenetic Modifications in Sperm: Unraveling Paternal Influences on Embryo Development and Transgenerational Health. Antioxidants (Basel). 2025 May 9;14(5):570.

27. Ntamo Y, Ziqubu K, Chellan N, Nkambule BB, Nyambuya TM, Mazibuko-Mbeje SE, et al. Drug-Induced Liver Injury: Clinical Evidence of N-Acetyl Cysteine Protective Effects. Oxid Med Cell Longev. 2021;2021:3320325.

28. Ahmadi S, Bashiri R, Ghadiri-Anari A, Nadjarzadeh A. Antioxidant supplements and semen parameters: An evidence based review. Int J Reprod Biomed. 2016 Dec;14(12):729–36.

29. Buhling K, Schumacher A, Eulenburg CZ, Laakmann E. Influence of oral vitamin and mineral supplementation on male infertility: a meta-analysis and systematic review. Reprod Biomed Online. 2019 Aug;39(2):269–79.

30. de Ligny W, Smits RM, Mackenzie-Proctor R, Jordan V, Fleischer K, de Bruin JP, et al. Antioxidants for male subfertility. Cochrane Database Syst Rev. 2022 May 4;5(5):CD007411.

31. Charron MJ, Williams L, Seki Y, Du XQ, Chaurasia B, Saghatelian A, et al. Antioxidant Effects of N-Acetylcysteine Prevent Programmed Metabolic Disease in Mice. Diabetes. 2020 Aug;69(8):1650–61.

32. Parnell SE, Sulik KK, Dehart DB, Chen S yu. Reduction of ethanol-induced ocular abnormalities in mice through dietary administration of N-acetylcysteine. Alcohol. 2010;44(7–8):699–705.

33. Percie du Sert N, Hurst V, Ahluwalia A, Alam S, Avey MT, Baker M, et al. The ARRIVE guidelines 2.0: updated guidelines for reporting animal research. BMJ Open Sci. 2020 July 20;4(1):e100115.

34. Cait J, Cait A, Scott RW, Winder CB, Mason GJ. Conventional laboratory housing increases morbidity and mortality in research rodents: results of a meta-analysis. BMC Biol. 2022 Jan 13;20(1):15.

35. Marian AJ, Senthil V, Chen SN, Lombardi R. Antifibrotic effects of antioxidant N-acetylcysteine in a mouse model of human hypertrophic cardiomyopathy mutation. J Am Coll Cardiol. 2006 Feb 21;47(4):827–34.

36. Wright DJ, Renoir T, Smith ZM, Frazier AE, Francis PS, Thorburn DR, et al. N-Acetylcysteine improves mitochondrial function and ameliorates behavioral deficits in the R6/1 mouse model of Huntington’s disease. Transl Psychiatry. 2015 Jan;5(1):e492.

37. Hu X, Chandler JD, Orr ML, Hao L, Liu K, Uppal K, et al. Selenium Supplementation Alters Hepatic Energy and Fatty Acid Metabolism in Mice. J Nutr. 2018 May 1;148(5):675–84.

38. Adler ID. Comparison of the duration of spermatogenesis between male rodents and humans. Mutat Res. 1996 June 10;352(1–2):169–72.

39. Whitten WK, Bronson FH, Greenstein JA. Estrus-inducing pheromone of male mice: transport by movement of air. Science. 1968 Aug 9;161(3841):584–5.

40. Truett GE, Heeger P, Mynatt RL, Truett AA, Walker JA, Warman ML. Preparation of PCR-quality mouse genomic DNA with hot sodium hydroxide and tris (HotSHOT). Biotechniques. 2000 July;29(1):52, 54.

41. Lesciotto KM, Perrine SMM, Kawasaki M, Stecko T, Ryan TM, Kawasaki K, et al. Phosphotungstic acid enhanced microCT: optimized protocols for embryonic and early postnatal mice. Dev Dyn. 2020 Apr;249(4):573–85.

42. Clercq KD, Persoons E, Napso T, Luyten C, Parac-Vogt TN, Sferruzzi-Perri AN, et al. High-resolution contrast-enhanced microCT reveals the true three-dimensional morphology of the murine placenta. PNAS. 2019 July 9;116(28):13927–36.

43. Rohlf FJ. tpsDig, digitize landmarks and outlines, version 2.05. Department of Ecology and Evolution, State University of New York at Stony Brook. 2005;

44. Rohlf FJ. tpsUtil. file utility program–Department of Ecology and Evolution, State University of New York at Stony Brook. Search in. 2015;

45. Klingenberg CP. MorphoJ: an integrated software package for geometric morphometrics. Mol Ecol Resour. 2011 Mar;11(2):353–7.

46. Hammer O, Harper DAT, Ryan PD. PAST: Paleontological Statistics Software Package for Education and Data Analysis.

47. Zelditch ML, Swiderski DL, Sheets HD. Geometric Morphometrics for Biologists: A Primer. Academic Press; 2012. 489 p.

48. Attanasio C, Nord AS, Zhu Y, Blow MJ, Li Z, Liberton DK, et al. Fine tuning of craniofacial morphology by distant-acting enhancers. Science. 2013 Oct 25;342(6157):1241006.

49. Wiseman DN, Samra N, Román Lara MM, Penrice SC, Goddard AD. The Novel Application of Geometric Morphometrics with Principal Component Analysis to Existing G Protein-Coupled Receptor (GPCR) Structures. Pharmaceuticals. 2021 Sept 23;14(10):953.

50. Geometric Morphometrics Tutorial | Sam Penrice [Internet]. [cited 2023 Feb 9]. Available from: https://sampenrice.com/geometric-morphometrics-tutorial/

51. Amrhein V, Greenland S, McShane B. Scientists rise up against statistical significance. Nature. 2019 Mar;567(7748):305–7.

52. Muff S, Nilsen EB, O’Hara RB, Nater CR. Rewriting results sections in the language of evidence. Trends Ecol Evol. 2022 Mar;37(3):203–10.

53. Falach-Malik A, Rozenfeld H, Chetboun M, Rozenberg K, Elyasiyan U, Sampson SR, et al. N-Acetyl-L-Cysteine inhibits the development of glucose intolerance and hepatic steatosis in diabetes-prone mice. Am J Transl Res. 2016;8(9):3744–56.

54. Castellani CA, Longchamps RJ, Sun J, Guallar E, Arking DE. Thinking outside the nucleus: mitochondrial DNA copy number in health and disease. Mitochondrion. 2020 July;53:214–23.

55. Caro AA, Bell M, Ejiofor S, Zurcher G, Petersen DR, Ronis MJJ. N-acetylcysteine inhibits the upregulation of mitochondrial biogenesis genes in livers from rats fed ethanol chronically. Alcohol Clin Exp Res. 2014 Dec;38(12):2896–906.

56. Coan PM, Ferguson-Smith AC, Burton GJ. Developmental dynamics of the definitive mouse placenta assessed by stereology. Biol Reprod. 2004 June;70(6):1806–13.

57. Mu J, Slevin JC, Qu D, McCormick S, Adamson SL. In vivo quantification of embryonic and placental growth during gestation in mice using micro-ultrasound. Reprod Biol Endocrinol. 2008 Aug 12;6:34.

58. Kaminen-Ahola N, Ahola A, Maga M, Mallitt KA, Fahey P, Cox TC, et al. Maternal ethanol consumption alters the epigenotype and the phenotype of offspring in a mouse model. PLoS Genet. 2010;6(1):e1000811.

59. Klingenberg C, Wetherill L, Rogers J, Moore E, Ward R, Autti-Rämö I, et al. Prenatal Alcohol Exposure Alters the Patterns of Facial Asymmetry. Alcohol. 2010;44(7–8):649–57.

60. Mutsvangwa TEM, Meintjes EM, Viljoen DL, Douglas TS. Morphometric analysis and classification of the facial phenotype associated with fetal alcohol syndrome in 5- and 12-year-old children. Am J Med Genet A. 2010 Jan;152A(1):32–41.

61. Anthony B, Vinci-Booher S, Wetherill L, Ward R, Goodlett C, Zhou FC. Alcohol-induced facial dysmorphology in C57BL/6 mouse models of fetal alcohol spectrum disorder. Alcohol. 2010 Dec;44(7–8):659–71.

62. Zhong Z, Lemasters JJ. A Unifying Hypothesis Linking Hepatic Adaptations for Ethanol Metabolism to the Proinflammatory and Profibrotic Events of Alcoholic Liver Disease. Alcohol Clin Exp Res. 2018 Nov;42(11):2072–89.

63. Simon L, Molina PE. Cellular Bioenergetics: Experimental Evidence for Alcohol-induced Adaptations. Function (Oxf). 2022;3(5):zqac039.

64. Schwalfenberg GK. N-Acetylcysteine: A Review of Clinical Usefulness (an Old Drug with New Tricks). J Nutr Metab. 2021 June 9;2021:9949453.

65. Ngo V, Karunatilleke NC, Brickenden A, Choy WY, Duennwald ML. Oxidative Stress-Induced Misfolding and Inclusion Formation of Nrf2 and Keap1. Antioxidants (Basel). 2022 Jan 27;11(2):243.

66. Lizzo G, Migliavacca E, Lamers D, Frézal A, Corthesy J, Vinyes-Parès G, et al. A Randomized Controlled Clinical Trial in Healthy Older Adults to Determine Efficacy of Glycine and N-Acetylcysteine Supplementation on Glutathione Redox Status and Oxidative Damage. Front Aging. 2022;3:852569.

67. Paschalis V, Theodorou AA, Margaritelis NV, Kyparos A, Nikolaidis MG. N-acetylcysteine supplementation increases exercise performance and reduces oxidative stress only in individuals with low levels of glutathione. Free Radic Biol Med. 2018 Feb 1;115:288–97.

68. Skvarc DR, Dean OM, Byrne LK, Gray L, Lane S, Lewis M, et al. The effect of N-acetylcysteine (NAC) on human cognition - A systematic review. Neurosci Biobehav Rev. 2017 July;78:44–56.

69. Peris E, Micallef P, Paul A, Palsdottir V, Enejder A, Bauzá-Thorbrügge M, et al. Antioxidant treatment induces reductive stress associated with mitochondrial dysfunction in adipocytes. J Biol Chem. 2019 Feb 15;294(7):2340–52.

70. Argaev-Frenkel L, Rosenzweig T. Complexity of NAC Action as an Antidiabetic Agent: Opposing Effects of Oxidative and Reductive Stress on Insulin Secretion and Insulin Signaling. Int J Mol Sci. 2022 Mar 9;23(6):2965.

71. Merry TL, Ristow M. Do antioxidant supplements interfere with skeletal muscle adaptation to exercise training? J Physiol. 2016 Sept 15;594(18):5135–47.

72. Higgins MR, Izadi A, Kaviani M. Antioxidants and Exercise Performance: With a Focus on Vitamin E and C Supplementation. Int J Environ Res Public Health. 2020 Nov 15;17(22):8452.

73. Deenadayalan A, Subramanian V, Paramasivan V, Veeraraghavan VP, Rengasamy G, Coiambatore Sadagopan J, et al. Stevioside Attenuates Insulin Resistance in Skeletal Muscle by Facilitating IR/IRS-1/Akt/GLUT 4 Signaling Pathways: An In Vivo and In Silico Approach. Molecules. 2021 Dec 20;26(24):7689.

