## Supplemental Table 1 for "Therapy to Teratology: Chronic Paternal Antioxidant Supplementation Alters Offspring Placental Architecture and Craniofacial Morphogenesis in a Mouse Model"

PROCRUSTES ANOVA RESULTS

| View | Procrustes ANOVA p-value | Centroid p-value |
| --- | --- | --- |
| Male Front | p < 0.0001 | p = 0.0455 |
| Male Left | p < 0.0001 | p = 0.4496 |
| Male Right | p = 1.0000 | p = 0.7429 |
| Female Front | p < 0.0001 | p = 0.4857 |
| Female Left | p < 0.0001 | p = 0.0761 |
| Female Right | p = 1.0000 | p = 0.0305 |

MANOVA RESULTS, FRONT (Bonferroni Corrected)

|  | Control Female | Control Male | Antioxidant Female | Antioxidant Male |
| --- | --- | --- | --- | --- |
| Control Female |  | 2.9513E-08 | 5.8773E-14 | 1.1108E-14 |
| Control Male | 2.9513E-08 |  | 1.0199E-11 | 2.8126E-14 |
| Antioxidant Female | 5.8773E-14 | 1.0199E-11 |  | 3.0212E-15 |
| Antioxidant Male | 1.1108E-14 | 2.8126E-14 | 3.0212E-15 |  |

ONE-WAY ANOSIM, FRONT (Bonferroni Corrected)

|  | Control Female | Control Male | Antioxidant Female | Antioxidant Male |
| --- | --- | --- | --- | --- |
| Control Female |  | 0.0006 | 0.0006 | 0.0006 |
| Control Male | 0.0006 |  | 0.0006 | 0.0006 |
| Antioxidant Female | 0.0006 | 0.0006 |  | 0.0006 |
| Antioxidant Male | 0.0006 | 0.0006 | 0.0006 |  |

ONE-WAY PERMANOVA, FRONT (Bonferroni Corrected)

|  | Control Female | Control Male | Antioxidant Female | Antioxidant Male |
| --- | --- | --- | --- | --- |
| Control Female |  | 0.0006 | 0.0006 | 0.0006 |
| Control Male | 0.0006 |  | 0.0006 | 0.0006 |
| Antioxidant Female | 0.0006 | 0.0006 |  | 0.0006 |
| Antioxidant Male | 0.0006 | 0.0006 | 0.0006 |  |

MANOVA RESULTS, LEFT (Bonferroni Corrected)

|  | Control Female | Control Male | Antioxidant Female | Antioxidant Male |
| --- | --- | --- | --- | --- |
| Control Female |  | 1.1084E-11 | 7.3072E-13 | 6.1209E-16 |
| Control Male | 1.1084E-11 |  | 8.8924E-15 | 9.4514E-15 |
| Antioxidant Female | 7.3072E-13 | 8.8924E-15 |  | 2.7162E-13 |
| Antioxidant Male | 6.1209E-16 | 9.4514E-15 | 2.7162E-13 |  |

ONE-WAY ANOSIM, LEFT (Bonferroni Corrected)

|  | Control Female | Control Male | Antioxidant Female | Antioxidant Male |
| --- | --- | --- | --- | --- |
| Control Female |  | 0.0006 | 0.0006 | 0.0006 |
| Control Male | 0.0006 |  | 0.0006 | 0.0006 |
| Antioxidant Female | 0.0006 | 0.0006 |  | 0.0006 |
| Antioxidant Male | 0.0006 | 0.0006 | 0.0006 |  |

ONE-WAY PERMANOVA, LEFT (Bonferroni Corrected)

|  | Control Female | Control Male | Antioxidant Female | Antioxidant Male |
| --- | --- | --- | --- | --- |
| Control Female |  | 0.0006 | 0.0006 | 0.0006 |
| Control Male | 0.0006 |  | 0.0006 | 0.0006 |
| Antioxidant Female | 0.0006 | 0.0006 |  | 0.0006 |
| Antioxidant Male | 0.0006 | 0.0006 | 0.0006 |  |

MANOVA RESULTS, RIGHT (Bonferroni Corrected)

|  | Control Female | Control Male | Antioxidant Female | Antioxidant Male |
| --- | --- | --- | --- | --- |
| Control Female |  | 3.1295E-11 | 1.7522E-12 | 1.9128E-11 |
| Control Male | 3.1295E-11 |  | 9.3827E-14 | 2.2414E-12 |
| Antioxidant Female | 1.7522E-12 | 9.3827E-14 |  | 5.3528E-13 |
| Antioxidant Male | 1.9128E-11 | 2.2414E-12 | 5.3528E-13 |  |

ONE-WAY ANOSIM, RIGHT (Bonferroni Corrected)

|  | Control Female | Control Male | Antioxidant Female | Antioxidant Male |
| --- | --- | --- | --- | --- |
| Control Female |  | 0.0006 | 0.0006 | 0.0006 |
| Control Male | 0.0006 |  | 0.0006 | 0.0006 |
| Antioxidant Female | 0.0006 | 0.0006 |  | 0.0006 |
| Antioxidant Male | 0.0006 | 0.0006 | 0.0006 |  |

ONE-WAY PERMANOVA, RIGHT(Bonferroni Corrected)

|  | Control Female | Control Male | Antioxidant Female | Antioxidant Male |
| --- | --- | --- | --- | --- |
| Control Female |  | 0.0006 | 0.0006 | 0.0006 |
| Control Male | 0.0006 |  | 0.0006 | 0.0006 |
| Antioxidant Female | 0.0006 | 0.0006 |  | 0.0006 |
| Antioxidant Male | 0.0006 | 0.0006 | 0.0006 |  |
